# Nicotinic Modulation of Fast-spiking Neurons in Rat Somatosensory Cortex Across Development

**DOI:** 10.1101/2025.06.18.660362

**Authors:** Catherine W. Haga, Jeffrey Koenig, Nathan Cramer, Ramesh Chandra, Asaf Keller

## Abstract

Signaling at nicotinic acetylcholine receptors (nAChRs) is vital for normal development of cerebral cortical circuits. These developing circuits are also shaped by fast-spiking (FS) inhibitory cortical neurons. While nicotinic dysfunction in FS neurons is implicated in a number of psychiatric and neurodevelopmental disorders, FS neurons are thought to not have nicotinic responses in adults. Here, we establish a timeline of FS neuron response to nicotine pre- and postsynaptically in primary somatosensory cortex in male and female rats. We found that nicotine increases the frequency of spontaneous synaptic inputs to FS neurons during the second postnatal week, and this effect persisted through development. In contrast, FS neurons in S1 had no postsynaptic responses to nicotine from as early as they can be reliably identified. This was not attributable to receptor desensitization, and we further revealed that FS neurons express abundant mRNA for several nAChR subunits, beginning early in development. To determine why FS neurons do not respond to nicotine, despite expressing these receptors, we probed for the expression of lynx1, a negative nicotinic modulator. Lynx1 mRNA was expressed in FS neurons from early development, with expression increasing dramatically during the second postnatal week.

**SIGNIFICANCE STATEMENT:** Signaling at nicotinic receptors is critical for development of cortical circuits. These circuits are also shaped by fast-spiking (FS) inhibitory neurons. We reveal how these developmental processes interact, by establishing a timeline of nicotinic effects on FS neurons in rats. We find that nicotine presynaptically regulates inputs to FS neurons from early development. However, FS neurons at all ages lack postsynaptic responses to nicotine, despite expressing nicotinic receptor mRNA. This might be due to the expression of lynx1, a negative nicotinic receptor modulator, which we identify in FS neurons as early as the first postnatal week. This work reveals novel aspects of development, relevant to both normal cortical development and the neuropsychiatric pathologies associated with abnormal FS neurons and nicotinic development.

## INTRODUCTION

Acetylcholine is the endogenous neurotransmitter that acts on muscarinic and nicotinic acetylcholine receptors (nAChRs), which are expressed widely in neocortex and are important for cortical development (Fiedler et al., 1987; Dwyer et al., 2008). Nicotinic acetylcholine receptors are upregulated during early development and their expression in the cortex peaks by the end of the second postnatal week (Fiedler et al., 1987; Naeff et al., 1992; Tribollet et al., 2004). Development is also driven by fast-spiking, parvalbumin-expressing GABAergic neurons (FS or PV neurons), a large class of inhibitory cortical neurons with characteristic kinetic properties (Hensch, 2005). During the first postnatal weeks, the biophysical and chemical properties of FS neurons mature (Itami et al., 2007). FS neurons drive use-dependent plasticity and circuit development during developmental critical periods (Hensch, 2005; Lee et al., 2012; Choi, 2018; Rupert & Shea, 2022; Kaplan et al., 2016).

Nicotinic signaling in FS neurons is implicated in neuropsychiatric disorders that involve network dysfunction (Lin et al., 2014; Deutsch & Burket, 2020). Mutation or microdeletion of the CHRNA7 gene coding for the α7 homomeric receptor is associated with schizophrenia, intellectual disability, autism spectrum disorder, and epilepsy (Sharp et al., 2008; Lin et al., 2014; Tregellas & Wylie, 2019). Behavioral abnormalities in attention, working memory, and learning associated with these disorders have also been functionally linked to deletion of this receptor in rodent models (Lin et al., 2014; Deutsch & Burkett, 2020). FS neurons are implicated in the mechanisms of these associations, as deletion of the α7 gene causes abnormal FS neuron development (Lin et al., 2014). Thus, nicotinic signaling in FS neurons is critical for normal cortical development.

There is some debate as to whether PV neurons express functional nAChRs. While some studies report that PV neurons may not express nAChRs in rodents (Couey et al., 2007; Askew et al., 2019), co-expression of both α7 and α4β2 nAChRs in PV neurons has been demonstrated in human temporal cortex (Krenz et al., 2001), and expression of β2 subunits has been identified in PV neurons in macaque visual cortex (Disney et al., 2007). However, FS neurons do not respond postsynaptically to nicotine in rodents, at least during late postnatal development through adulthood. This has been confirmed in auditory, visual, somatosensory, and prefrontal cortical areas (Couey et al., 2007; Gulledge et al., 2007; Askew et al., 2019). In contrast, whether FS neurons respond to nicotine at earlier developmental ages has not been established. Here, we address this deficiency by studying the actions of nicotine on both presynaptic and postsynaptic responses of FS neurons across development.

We establish the developmental timeline of FS neurons’ responses to nicotine in the barrel field of primary somatosensory cortex (S1), a well-established developmental model of cortical circuits and functions (Erzurumlu & Gaspar, 2012; Erzurumlu & Gaspar, 2020). We also investigate the expression of lynx1, an endogenous prototoxin that serves as a brake on nicotinic signaling in other cortical areas (Morishita et al., 2010), with a focus on its specific expression in fast-spiking (FS) interneurons in S1 across postnatal development.

## MATERIALS AND METHODS

### Animals

All procedures adhered to the Guide for the Care and Use of Laboratory Animals and were approved by the Institutional Animal Care and Use Committee at the University of Maryland School of Medicine. Male and female Long-Evans rats from Charles River were bred in our temperature- and humidity-controlled vivarium. Animals were fed standard chow *ad libitum* and maintained on a 12-hour light/dark cycle. Rats were bred in monogamous pairs with gestational timing confirmed by the presence of sperm in vaginal lavage samples. Dams were pair-housed then single-housed immediately before delivery until study endpoint. Litters were culled to 8-12 pups per dam. Preweanling offspring were used in most experiments. In some cases, pups were weaned at postnatal day 21 (P21) and socially housed until study endpoint. Preweanling (P12) and adult (P100) male and female C57BL/6 mice were used for lynx1 RNAscope™ experiments.

### Slice electrophysiology

In vitro slice electrophysiology was performed as we previously described (Alipio et al., 2021). We anesthetized preweanling and adolescent rats with ketamine (40-80 mg/kg) and xylazine (5-10 mg/kg), removed their brains, and prepared 300-μm coronal sections containing primary somatosensory cortex using a Leica VT1200s vibratome. Slices from preweanling rats up to P14 were prepared in ice cold normal artificial cerebrospinal fluid (ACSF) containing the following (in mM): 119 NaCl, 2.5 KCl, 1.2 NaH2PO4, 2.4 NaHCO3, 12.5 glucose, 2 MgSO4·7H2O, and 2 CaCl2·2H2O. Slices from rats P15 and older were prepared in ice cold NMDG ACSF containing the following (in mM): 92 NMDG, 30 sodium bicarbonate, 20 HEPES, 25 glucose, 5 sodium ascorbate, 2 thiourea, 1.25 monosodium phosphate, 2.5 potassium chloride, 3 sodium pyruvate, 10 magnesium sulfate heptahydrate, and 0.5 calcium chloride dihydrate. Slices prepared in NMDG ACSF were allowed to recover in 35–37°C NMDG ACSF for 7 minutes immediately after slicing before being placed in room temperature normal ACSF. All ACSF solutions were adjusted to a pH of 7.4 using HCl, and osmolarity was adjusted to 305 ± 5 mOsm/L. Solutions were saturated with carbogen (95% O2 and 5% CO2) throughout use.

We placed slices in a submersion chamber and continually perfused (2 ml/min) with normal ACSF. We obtained whole-cell patch-clamp recordings from S1 layer 4 (L4) neurons first in current-clamp mode to establish their kinetic parameters from intracellular current injections. Using a series of 1000 ms current pulses of increasing amplitude, we recorded and analyzed the firing pattern of each neuron for the first 500 ms of stimulation. We subsequently recorded spontaneous postsynaptic currents (sPSCs) and measurements of whole cell resistance in voltage-clamp mode (-65 mV holding potential) through pipettes (4-6 MΩ) containing (in mM): 120 potassium gluconate, 10 potassium chloride, 10 HEPES, 1 magnesium chloride, 0.5 EGTA, 2.5 magnesium ATP, 0.2 GTP-Tris, and 0.1% biocytin (Thermo Fisher Scientific), adjusted to pH 7.3 and 290 mOsm/L. Biocytin was included in the internal solution to allow for reconstruction and further identification of recorded cells. In some sPSC experiments gabazine (1 μM) was added to the ACSF to suppress the effects of inhibitory inputs on the recorded cells acting on GABA-A receptors and isolate excitatory inputs. However, even in cases where we did not block GABA-A signaling, most inhibitory currents driven by chloride would not be detectable due to a lack of driving force for this ion. For recordings in voltage clamp mode, cells were held at -65 mV— effectively the reversal potential of chloride with our solutions— and thus we assume that recorded sPSCs are glutamatergic in nature and refer to them in Results as sEPSCs. Nicotine bitartrate dihydrate (Sigma-Aldrich, St. Louis, MO) was applied in slice electrophysiology experiments by washing in with the ACSF for 3-6 minutes (10µM) or directly applying approximately 50 μm from the soma of recorded neurons through a glass pipette (4-6 MΩ), using a Picospritzer (20 to 30 µM).

We recorded baseline sEPSCs from FS neurons for a minimum of 3 minutes before washing in ACSF containing nicotine (10 μM) for 6 minutes. We compared frequency and amplitude of sEPSCs during 3 minutes of baseline and 3 minutes of nicotine application, following the initial 3-minute wash-in period. We recorded neuronal membrane resistance, an indirect measure of ionic currents, by measuring the voltage response during the steady-state period of intracellular current injections, before and after bath nicotine application for 3 minutes (10 μM).

In all current-clamp experiments, gabazine (1 μM), CNQX (20 μM), and AP5 (50 µM) were added to the ACSF to suppress synaptic activity and study somatodendritic effects of nicotine. We measured resting membrane potential and rheobase of neurons from P10-13 rats in current clamp configuration. These measurements were repeated after 3 minutes of bath nicotine application (10μM). Rheobase was calculated from 500 ms current steps in increments of 10-20 pA until cells reached action potential threshold. Rheobase was further refined for each neuron in 500 ms current steps in increments of 2 pA until a more precise threshold was reached.

For puff recordings, the puff pipette was attached to a Sutter Instruments micromanipulator for positioning and pressure ejection of nicotine solution controlled with a Picospritzer. Prior to recording, the puff pipette was lowered into a region of the brain outside of the cortex and the air pressure adjusted (∼ 5 to 10 PSI) so that the pressure wave extended at least 50 µm from the tip of the pipette while causing minimal mechanical disturbance to the cells at this range. A putative FS neuron was identified, and the puff pipette placed ∼ 50 µm away, but aimed directly at, the cell. A second pipette was used to obtain whole cell recordings. We checked for responses to nicotine puff application by comparing the area under the curve (AUC) of each recording 3 seconds prior to each puff application (baseline) and 1 second after the onset of puff application. The AUCs for the first 3 to 5 consecutive applications in each cell were compared using a paired *t*-test with *p* < 0.05 indicating a significant response to nicotine. For cells that showed a significant response to nicotine, we report the peak change in membrane potential. Cells that did not respond to nicotine were assigned a value of 0 mV.

Series resistance was monitored in electrophysiological recordings by measuring the current evoked by a -5 mV square pulse, and recordings were discarded if resistance changed by >20%. Recordings were made with the Axon pClamp 11 Software Suite (Molecular Devices, Silicon Valley, CA) and analyzed with Easy Electrophysiology V2 software (Easy Electrophysiology Ltd, London, England).

### Histology

After electrophysiological recordings, slices were fixed in 10% neutral buffered formalin (NBF) at room temperature for 24-48 hours and stored at 4° C in PBS until processing. Slices were placed in blocking solution containing PBS, 1% BSA, and 0.3% Triton for two hours, then incubated for 48-72 hours in incubation solution (1% BSA, 0.1% Triton) containing streptavidin-conjugated Alexa 488 (1:1000; Jackson ImmunoResearch Laboratories, West Grove, PA) and parvalbumin polyclonal antibody (1:10,000; PA1-933; Thermo Fisher Scientific). Slices were washed and incubated in a secondary antibody solution (Alexa Fluor 594 donkey anti-rabbit IgG, 1:500; Thermo Fisher Scientific) then mounted in aqueous media and imaged using confocal microscopy. Neurons were analyzed for parvalbumin immunoreactivity, soma and dendritic morphology, and presence of dendritic spines to confirm phenotype as FS or regular spiking (RS).

### RNAscope™

In situ mRNA expression levels were determined using RNAscope™ Multiplex Fluorescent V2 Assays (ACDBio, Newark, CA). Animals were deeply anesthetized by intraperitoneal injection of ketamine/xylazine and transcardially perfused with ice-cold PBS followed by 10% neutral buffered formalin (NBF). Brains were extracted, fixed overnight in 10% NBF, cryoprotected in a sucrose solution, frozen, and sectioned using a cryostat (12 µM). Ready-to-use reagents from the RNAscope™ Multiplex Fluorescent V2 Assay Kit were used with Rn-Chrna4, Rn-Chrna7, and Rn-Pvalb probes (ACDBio) to process sections from P12 and P19 male and female rats to detect mRNA for nAChR subunits and parvalbumin. Mm-Lynx1 and Mm-Pvalb probes (ACDBio) were used to process sections from P12 and P100 male and female mice. Sections were mounted in aqueous media and imaged using confocal microscopy. In situ mRNA expression levels were quantified using Imaris 10 software (Oxford Instruments, Abingdon, United Kingdom).

### RT-qPCR

Male and female rats were euthanized at P0, P4, P7, P14, and P21. Brains were extracted and dissected in ice-cold PBS to collect 14-gauge tissue punches from 1 mm sections of S1 and primary visual cortex (V1). Two brains were pooled for each region from P0 and P4 animals due to tissue volume. RNA was isolated from cortical tissue punches with Trizol reagent (Invitrogen) and the MicroElute Total RNA Kit (Omega; Cat# R6831) with a DNase step (Qiagen, Germantown, MD; Cat# 79254). RNA concentration was measured on a Nanodrop (Thermo Scientific) and 1000 ng of mRNA from each sample was used to synthesize complementary DNA using an iScript cDNA synthesis kit (Bio-Rad, Hercules, CA; Cat# 1708891). This cDNA was diluted to a concentration of 5 ng/μL, which was used to measure relative mRNA expression of lynx1 by age via quantitative PCR with PerfeCTa SYBR Green FastMix (Quantabio, Beverly, MA; Cat# 95072). Primer sets used were as follows (F, R; 5’-3’): Lynx1 ACCACTCGAACTTACTTCACC, ATCGTACACGGTCTCAAAGC; GAPDH CCCACTCTTCCACCTTCGATG, TCCACCACCCTGTTGCTGTAG. All samples were run in duplicate, and samples with a difference in CT value greater than 0.6 were excluded. Relative quantification of mRNA expression was performed using the 2^−ΔΔCt^ method with standard protocols, using Gapdh and P21 values in each respective cortical region to normalize expression.

### Statistical Analysis

Statistical tests were conducted using Graphpad Prism 10 software (Boston, MA). Sample sizes were determined a priori using G*Power software suite (Faul et al., 2007). Statistical significance level was set at *p* < 0.05. Effects of nicotine on individual neurons as well as group effects were analyzed. Inter-event intervals (IEI) for each sEPSC were calculated, and frequency is reported as the instantaneous frequency of events, or the inverse of IEIs. Effects of nicotine on frequency and amplitude of events from each neuron were separately analyzed (see Tables 1-2). We also calculated an average frequency and amplitude at baseline and with nicotine for each neuron and analyzed group effects with paired *t*-tests. We tested for sex differences in each experimental group where powered and grouped animals according to age if none were found. Where relevant, figures depict sex of the animal for each associated data point. Parametric tests were used when assumptions of normality were met, otherwise, nonparametric tests were used.

**Table 1.**
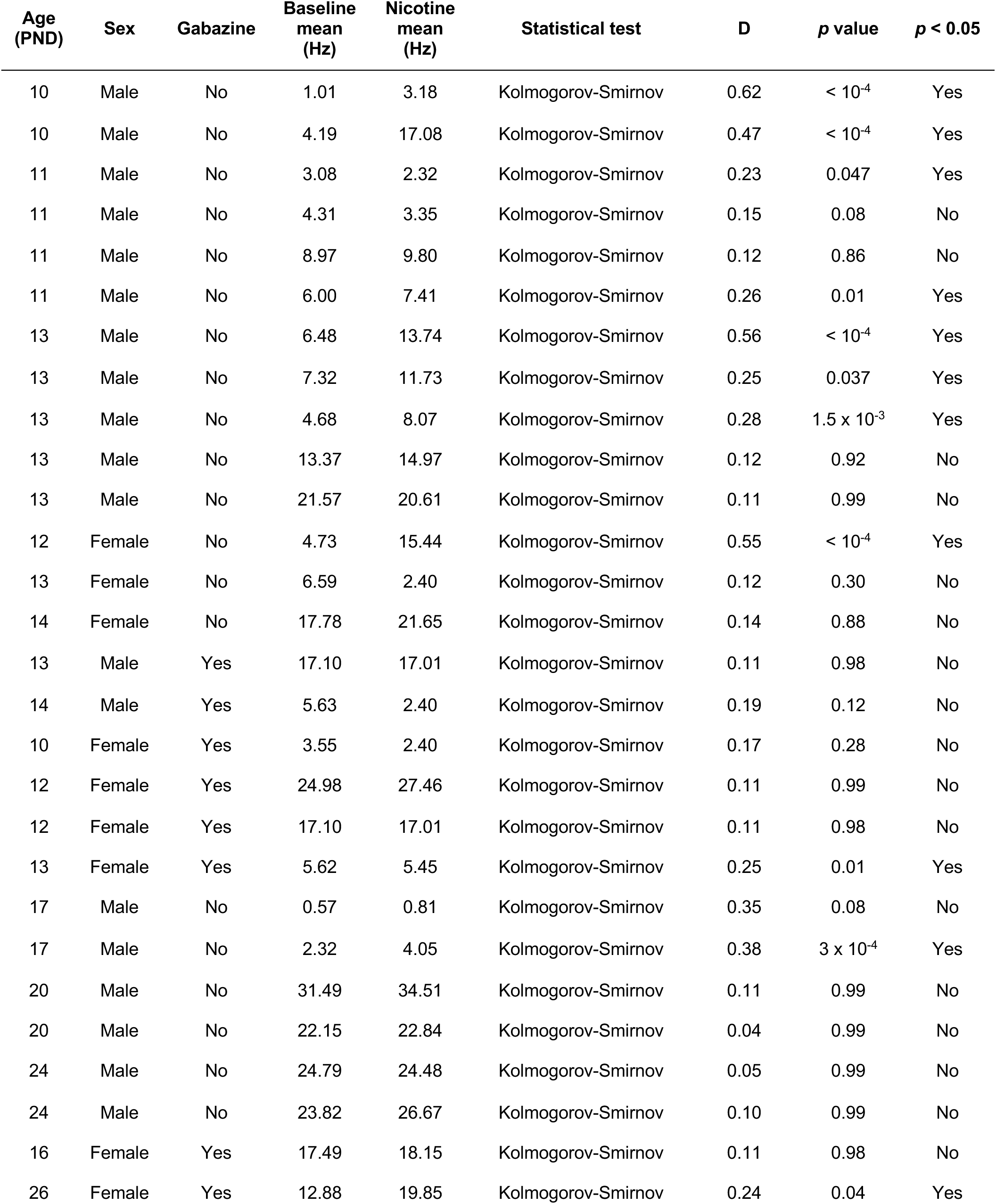
Frequency of sEPSCs by neuron.

| Age (PND) | Sex | Gabazine | Baseline mean (Hz) | Nicotine mean (Hz) | Statistical test | D | p value | p < 0.05 |
| --- | --- | --- | --- | --- | --- | --- | --- | --- |
| 10 | Male | No | 1.01 | 3.18 | Kolmogorov-Smirnov | 0.62 | < 10<sup>-4</sup> | Yes |
| 10 | Male | No | 4.19 | 17.08 | Kolmogorov-Smirnov | 0.47 | < 10<sup>-4</sup> | Yes |
| 11 | Male | No | 3.08 | 2.32 | Kolmogorov-Smirnov | 0.23 | 0.047 | Yes |
| 11 | Male | No | 4.31 | 3.35 | Kolmogorov-Smirnov | 0.15 | 0.08 | No |
| 11 | Male | No | 8.97 | 9.80 | Kolmogorov-Smirnov | 0.12 | 0.86 | No |
| 11 | Male | No | 6.00 | 7.41 | Kolmogorov-Smirnov | 0.26 | 0.01 | Yes |
| 13 | Male | No | 6.48 | 13.74 | Kolmogorov-Smirnov | 0.56 | < 10<sup>-4</sup> | Yes |
| 13 | Male | No | 7.32 | 11.73 | Kolmogorov-Smirnov | 0.25 | 0.037 | Yes |
| 13 | Male | No | 4.68 | 8.07 | Kolmogorov-Smirnov | 0.28 | 1.5 x 10<sup>-3</sup> | Yes |
| 13 | Male | No | 13.37 | 14.97 | Kolmogorov-Smirnov | 0.12 | 0.92 | No |
| 13 | Male | No | 21.57 | 20.61 | Kolmogorov-Smirnov | 0.11 | 0.99 | No |
| 12 | Female | No | 4.73 | 15.44 | Kolmogorov-Smirnov | 0.55 | < 10<sup>-4</sup> | Yes |
| 13 | Female | No | 6.59 | 2.40 | Kolmogorov-Smirnov | 0.12 | 0.30 | No |
| 14 | Female | No | 17.78 | 21.65 | Kolmogorov-Smirnov | 0.14 | 0.88 | No |
| 13 | Male | Yes | 17.10 | 17.01 | Kolmogorov-Smirnov | 0.11 | 0.98 | No |
| 14 | Male | Yes | 5.63 | 2.40 | Kolmogorov-Smirnov | 0.19 | 0.12 | No |
| 10 | Female | Yes | 3.55 | 2.40 | Kolmogorov-Smirnov | 0.17 | 0.28 | No |
| 12 | Female | Yes | 24.98 | 27.46 | Kolmogorov-Smirnov | 0.11 | 0.99 | No |
| 12 | Female | Yes | 17.10 | 17.01 | Kolmogorov-Smirnov | 0.11 | 0.98 | No |
| 13 | Female | Yes | 5.62 | 5.45 | Kolmogorov-Smirnov | 0.25 | 0.01 | Yes |
| 17 | Male | No | 0.57 | 0.81 | Kolmogorov-Smirnov | 0.35 | 0.08 | No |
| 17 | Male | No | 2.32 | 4.05 | Kolmogorov-Smirnov | 0.38 | 3 x 10<sup>-4</sup> | Yes |
| 20 | Male | No | 31.49 | 34.51 | Kolmogorov-Smirnov | 0.11 | 0.99 | No |
| 20 | Male | No | 22.15 | 22.84 | Kolmogorov-Smirnov | 0.04 | 0.99 | No |
| 24 | Male | No | 24.79 | 24.48 | Kolmogorov-Smirnov | 0.05 | 0.99 | No |
| 24 | Male | No | 23.82 | 26.67 | Kolmogorov-Smirnov | 0.10 | 0.99 | No |
| 16 | Female | Yes | 17.49 | 18.15 | Kolmogorov-Smirnov | 0.11 | 0.98 | No |
| 26 | Female | Yes | 12.88 | 19.85 | Kolmogorov-Smirnov | 0.24 | 0.04 | Yes |

**Table 2.** Amplitude of sEPSCs by neuron.

| Age (PND) | Sex | Gabazine | Baseline amplitude (pA) | Nicotine amplitude (pA) | Statistical test | D | p value | p < 0.05 |
| --- | --- | --- | --- | --- | --- | --- | --- | --- |
| 10 | Male | No | 17.75 | 18.42 | Kolmogorov-Smirnov | 0.18 | 0.79 | No |
| 10 | Male | No | 33.80 | 31.16 | Kolmogorov-Smirnov | 0.17 | 0.99 | No |
| 11 | Male | No | 17.40 | 17.70 | Kolmogorov-Smirnov | 0.11 | 0.95 | No |
| 11 | Male | No | 31.52 | 26.21 | Kolmogorov-Smirnov | 0.14 | 0.99 | No |
| 13 | Male | No | 11.87 | 10.96 | Kolmogorov-Smirnov | 0.53 | < 10<sup>-4</sup> | Yes |
| 13 | Male | No | 22.24 | 19.62 | Kolmogorov-Smirnov | 0.24 | 0.59 | No |
| 13 | Male | No | 22.35 | 18.76 | Kolmogorov-Smirnov | 0.26 | 0.53 | No |
| 13 | Male | No | 14.88 | 15.33 | Kolmogorov-Smirnov | 0.08 | 0.96 | No |
| 13 | Male | No | 25.98 | 33.85 | Kolmogorov-Smirnov | 0.22 | 0.03 | Yes |
| 12 | Female | No | 13.93 | 33.36 | Kolmogorov-Smirnov | 0.64 | < 10<sup>-4</sup> | Yes |
| 13 | Female | No | 22.31 | 15.55 | Kolmogorov-Smirnov | 0.23 | 2.6 x 10<sup>-3</sup> | Yes |
| 14 | Female | No | 22.41 | 23.01 | Kolmogorov-Smirnov | 0.15 | 0.75 | No |
| 13 | Male | Yes | 21.61 | 21.19 | Kolmogorov-Smirnov | 0.37 | 3.7 x 10<sup>-3</sup> | Yes |
| 14 | Male | Yes | 18.90 | 19.93 | Kolmogorov-Smirnov | 0.28 | 4.0 x 10<sup>-3</sup> | Yes |
| 10 | Female | Yes | 19.93 | 19.93 | Kolmogorov-Smirnov | 0.26 | 0.03 | Yes |
| 12 | Female | Yes | 20.97 | 20.62 | Kolmogorov-Smirnov | 0.13 | 0.73 | No |
| 12 | Female | Yes | 21.61 | 21.19 | Kolmogorov-Smirnov | 0.37 | 3.7 x 10<sup>-3</sup> | Yes |
| 13 | Female | Yes | 18.17 | 17.53 | Kolmogorov-Smirnov | 0.07 | 0.99 | No |
| 17 | Male | No | 13.57 | 14.66 | Kolmogorov-Smirnov | 0.27 | 0.28 | No |
| 17 | Male | No | 12.20 | 12.56 | Kolmogorov-Smirnov | 0.17 | 0.32 | No |
| 20 | Male | No | 22.39 | 21.21 | Kolmogorov-Smirnov | 0.20 | 0.17 | No |
| 20 | Male | No | 17.34 | 17.63 | Kolmogorov-Smirnov | 0.12 | 0.67 | No |
| 24 | Male | No | 22.68 | 22.94 | Kolmogorov-Smirnov | 0.18 | 0.66 | No |
| 24 | Male | No | 19.43 | 19.64 | Kolmogorov-Smirnov | 0.13 | 0.84 | No |
| 16 | Female | Yes | 21.68 | 21.46 | Kolmogorov-Smirnov | 0.18 | 0.41 | No |

## RESULTS

### Identifying PV Neurons

Fig. 1A depicts an FS neuron as viewed through diffusion interference contract (DIC) during live-slice electrophysiology with characteristic morphology including oblong shape and absence of a pronounced apical dendrite. FS neurons can be identified by the distinct properties of their action potentials, evoked by intracellular current injections (McCormick et al., 1985; Tateno et al., 2004; Subkhankulova et al., 2010). These include a brief spike duration, high firing frequency, a large afterhyperpolarization (AHP) (Fig. 1B), and little or no spike frequency adaptation (Fig 1C; 1D, left). Whenever possible, we confirmed the identity of these neurons with post hoc parvalbumin immunohistochemistry and verification of absence of dendritic spines (Fig. 1E). The FS characteristics contrast with those of regular spiking (RS) neurons that have longer spike durations, small or no AHP, and whose spike trains accommodate (Fig. 1D, right). RS cortical neurons may include both excitatory neurons and inhibitory neurons that are not FS and do not express PV (Porter et al., 1999; Rudy et al., 2011).

**Figure 1.**
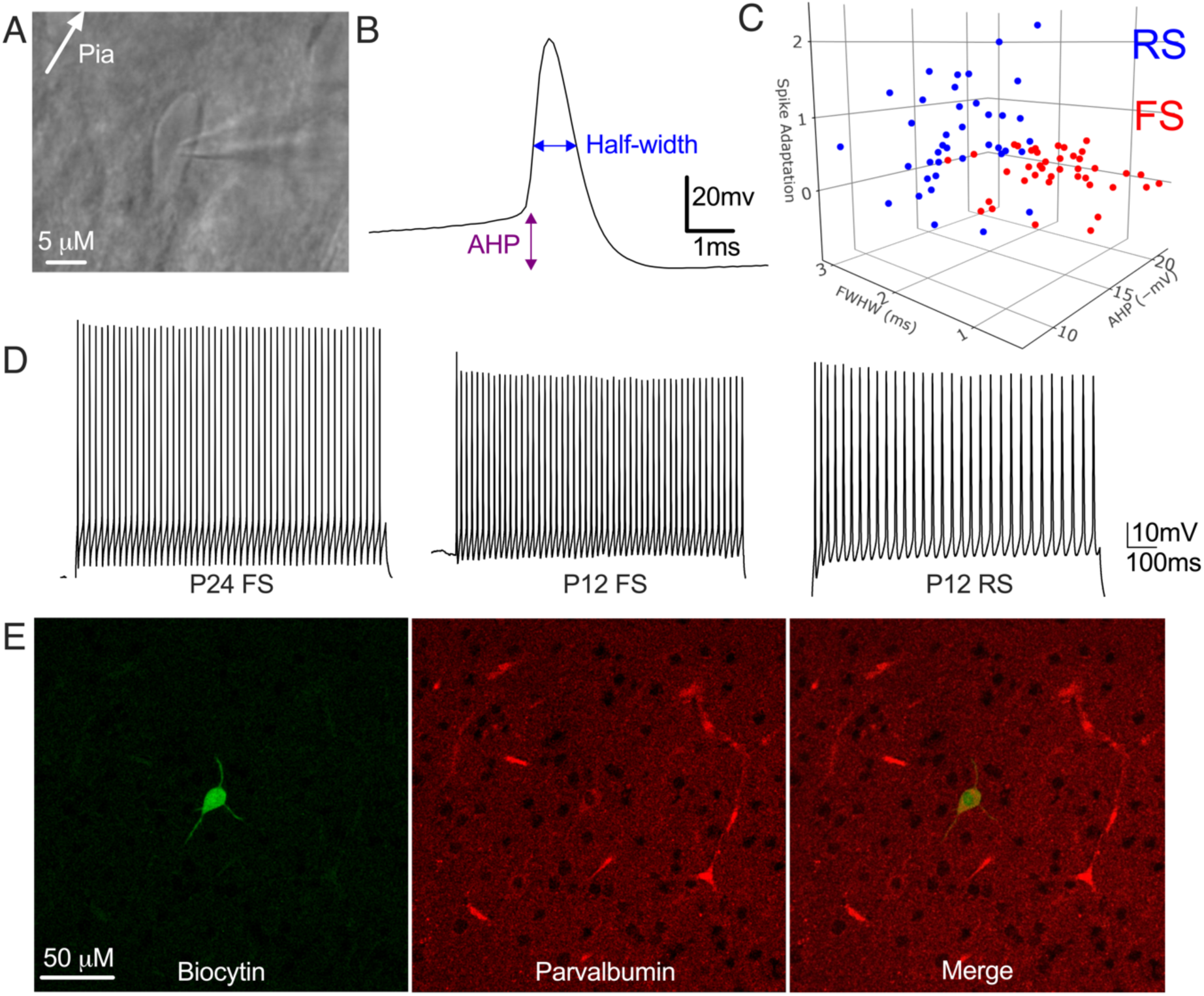
Identification of PV neurons in early development. **A**. FS neuron imaged using differential interference contrast during whole-cell patch clamp recording. **B**. Recorded action potential from an FS neuron. Spike half-width and AHP measurement locations indicated. **C**. Distribution of kinetic parameters used to identify neurons as FS or RS (Spike adaptation index, FWHW = full width at half maximum, or half-width, and AHP). Spike adaptation index is transformed by a scaling factor for simplicity of presentation. **D**. *Left:* Trace of action potentials recorded from an FS neuron from an adult rat (AHP: -21.7mV, HW: 0.6ms, IEI slope: 1.0e-5). Additional traces from neurons from P12 rats classified as FS (*center*; AHP: -17.9mV, HW: 1.5ms, IEI slope: 1.7e-4) or RS (*right*; AHP: -14.9mV, HW: 1.8ms, IEI slope: 1.0e-3). **E**. Biocytin-filled recorded neuron positive for parvalbumin IHC in a 300μm section.

It is more challenging to identify FS neurons in younger animals in which currents that drive the distinctive firing characteristics are still developing (Okaty et al., 2009; Subkhankulova et al., 2010), and in which significant PV expression begins only around P12 (Alcántara et al.,1993; De Lecea et al., 1995; Itami et al., 2007). To address this, we developed quantitative criteria to distinguish between FS and RS neurons. These criteria were obtained from recordings of 87 neurons from animals ranging in age from P7 to P26 (Fig. 1C) and were validated by IHC in a subset of cells (Fig. 1E). Neurons were identified as FS if they had AHP ≤ -10 mV, action potential width (at half maximum) ≤ 1.7 ms, and slope of inter-spike intervals (IEI slope, or spike adaptation index) ≤ 7.1e-4. Neurons that did not meet these criteria were classified as RS neurons. Fig. 1D depicts the differences in action potential kinetics obtained from P12 neurons classified as FS and RS, respectively. The depicted P12 FS neuron has an AHP of - 17.9 mV, half-width of 1.5 ms, and IEI slope of 1.7e-4, while the RS neuron has a longer half-width (1.8 ms) and more pronounced spike adaptation (IEI slope = 1.0e-3).

### Nicotine increases excitatory synaptic inputs to FS neurons in S1 across development

nAChRs are expressed by inhibitory, GABAergic cortical neurons, and by the presynaptic, glutamatergic terminals that provide excitatory inputs to these neurons (Gil et al., 1997; Demars & Morishita, 2014; Askew et al., 2019; Gil & Metherate, 2019). Previous studies examined the effects of nicotine on FS neurons after the second postnatal week and into adulthood. Therefore, we focused on comparing the second postnatal week (P8-P14) and the third week and beyond (P15-26). The second postnatal week in rats coincides with biophysical and chemical maturation of FS neurons (Itami et al., 2007) and a number of other important developmental changes. We investigated whether nAChR activation would increase synaptic drive onto FS neurons in early postnatal development and compared this to responses in early adolescence.

We recorded sEPSCs in S1 layer 4 FS neurons from postnatal rats (*N* = 19 rats P10-26, *n* = 1-2 neurons per rat). We assessed the frequency and amplitude of events from 3-minute recordings at baseline in ACSF and after a 3-minute wash-in of 10 µM nicotine. Frequency and amplitude of events were increased in some neurons by application of nicotine. Fig. 2A depicts traces of sEPSCs before and after nicotine application recorded from a FS neuron that significantly increased frequency and amplitude during the second postnatal week. A representative cumulative probability distribution curve for a neuron that responded to nicotine with an increased frequency of sEPSCs is shown in Fig. 2B, and for a neuron that responded to nicotine with increased amplitude of sEPSCs is shown in Fig. 2C. Nicotine produced significant rightward shifts in the cumulative probability distribution of frequency and amplitude, respectively, of each neuron depicted. Most (57%) neurons in the younger age group showed an increase in frequency (Fig. 2D, top left), and 25% exhibited an increase in sEPSC amplitude (Fig. 2E, top left). In 7% of recorded FS neurons in younger animals, frequency and amplitude increased (see Tables 1-2 for individual statistical tests). Nicotine significantly increased the mean frequency of sEPSCs in group analysis (Fig. 2D, bottom left), but not amplitude (Fig. 2E, bottom left).

**Figure 2.**
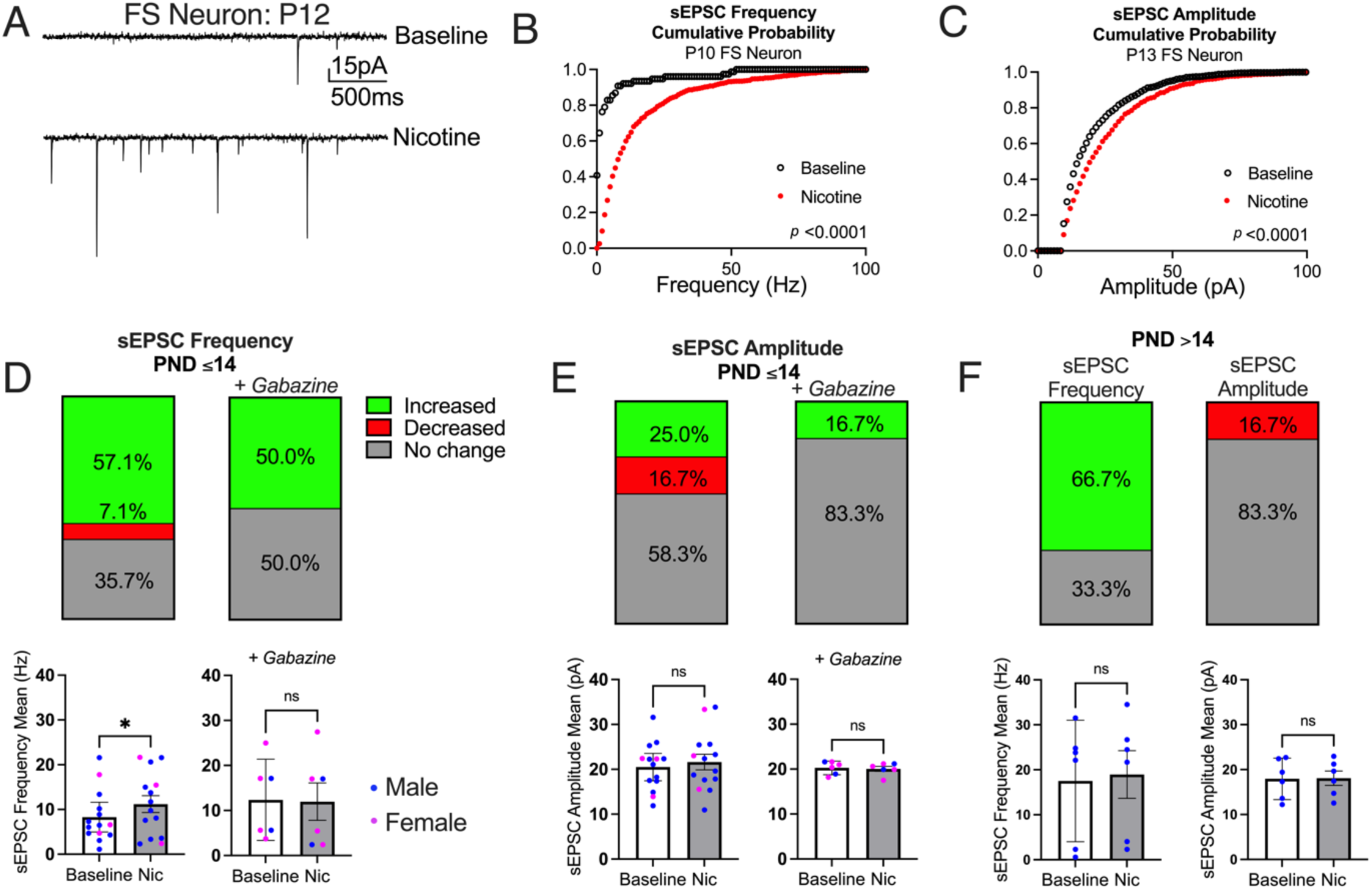
Nicotine enhances excitatory inputs to FS neurons. **A**. sEPSCs recorded from a P12 FS neuron that responded to nicotine by increasing frequency and amplitude of sEPSCs. **B**. Cumulative probability distribution of sEPSC frequency from a P10 FS neuron (Mann Whitney test, *U* = 6,588, *p* < 0.0001). **C**. Cumulative probability distribution of sEPSC amplitude from a P13 FS neuron (Mann Whitney test, *U* = 669,189, *p* < 0.0001). Percentages of FS neurons from P10-14 animals that significantly increased frequency (**D,** *n* = 14 neurons without gabazine; *n* = 6 neurons with gabazine; see Table 1 for individual statistical tests) and amplitude (**E,** *n* = 12 neurons without gabazine; *n* = 6 neurons with gabazine; see Table 2 for individual statistical tests) of sEPSCs with nicotine. Individual means of sEPSC frequency after nicotine application in the absence of (paired *t*-test, t(13) = 2.44, *p* = 0.03, Cohen’s *d* = 0.65) and presence of (paired *t*-test, t_(5)_ = 0.53, *p* = 0.62) gabazine are shown in **D**, lower panels. Individual means of sEPSC amplitude after nicotine application in the absence of (paired *t*-test, t_(13)_ = 0.64, *p* = 0.53) and presence of (paired *t*-test, t_(5)_ = 0.84, *p* = 0.44) gabazine are in **E**, lower panels. **F**. Percentages of FS neurons from P15-26 animals that significantly increased frequency or decreased amplitude of sEPSCs with nicotine (*n* = 6 neurons, see Tables 1-2 for individual statistical tests). Individual means of sEPSC frequency (*lower left*, paired *t*-test, t_(5)_ = 2.33, *p* = 0.07) and sEPSC amplitude (*lower right*, paired *t*-test, t_(5)_ = 0.49, *p* = 0.65). Means ± 95% CI (**D**, **E**, **F**, *lower panels*).

In a subset of recordings from FS neurons in P10-14 rats, we included the GABA-A receptor antagonist, gabazine (1 µM) in the ACSF to suppress effects of inhibitory interneurons on the recorded cells. In this condition, similar ratios of FS neurons responded to nicotine with an increased frequency (50%, Fig. 2D, top right) and amplitude (17%, Fig. 2E, top right). However, the group-level enhancement of frequency was abolished when gabazine was included in the recording solution (Fig. 2D, bottom right). Nicotine may partially be acting on inhibitory neurons to drive increases in sEPSC frequency in FS neurons at young ages. As some of these recordings occurred before GABA currents switch from excitatory to inhibitory, a developmental milestone that occurs by the end of the second postnatal week (Peerboom & Wierenga, 2021), the increased frequency from nicotine application at early ages may be directly driven by excitatory GABA signaling. We have also considered an indirect mechanism involving disinhibition via other interneurons, as has been proposed in other studies (Askew, et al., 2019).

Conversely, in older animals (P15-26), there were no group effects of nicotine on either the frequency (Fig. 2F, bottom left) or amplitude (Fig. 2F, bottom right) of sEPSCs. While a similar proportion of neurons in this older age group still showed increased sEPSC frequency (Fig. 2F, top left), none of the neurons responded with an increase in amplitude at these ages (Fig. 2F, top right). These results collectively indicate that while nicotine increases the frequency of excitatory synaptic events in a majority of FS neurons across development (from the second postnatal week into adolescence), the group-level impact on frequency is primarily evident in younger animals, and the amplitude-enhancing effects are specifically limited to this early developmental period in a smaller subset of FS neurons.

### Nicotine does not produce whole-cell currents in FS neurons in S1

We also explored direct postsynaptic effects of nicotine on FS neurons in S1, beginning in early development. While previous research in adult rodents has generally reported that fast-spiking (FS) neurons do not demonstrate direct postsynaptic nicotinic currents in various cortical regions in adults, including prefrontal cortex (Couey et al., 2007; Gulledge et al., 2007), motor cortex (Porter et al., 1999), visual cortex and somatosensory cortex (Gulledge et al., 2007), our study specifically investigates these effects in earlier developmental periods in S1. In contrast to FS neurons, regular-spiking (RS) neurons across multiple cortical areas typically depolarize in response to nicotine, whether applied via bath (Askew et al., 2019) or directly to the soma (Gulledge et al., 2007; Couey et al., 2007). Consequently, some have suggested that FS neurons may not possess functional nAChRs in rodents (Couey et al., 2007; Askew et al., 2019). There is some evidence from other species demonstrating expression of nAChR subunits in PV-expressing neurons (Krenz et al., 2001; Disney et al, 2007), though to our knowledge this has not been explored in rats.

We hypothesized that nicotine may have postsynaptic effects on FS neurons at very young ages, and again focused on specific differences in effects between the second postnatal week and after P15. We calculated normalized neuronal resistance changes, an indirect measure of whole-cell currents, after applying nicotine (10 µM) for 3 min. An example I-V curve used to produce these data is shown in Fig. 3A. Nicotine application did not decrease whole-cell resistance in FS neurons at any age range examined, from P8-26 (Fig. 3B, left), P8-14 (Fig. 3C, left), or P15-26 (Fig. 3F, left). In a subset of FS neurons from young (Fig. 3C, right) and older (Fig. 3D, right) animals, we recorded whole-cell resistance changes with nicotine in the presence of gabazine (1 µM) and similarly found no effect. To investigate effects of nicotine on intrinsic excitability of FS neurons at young ages, we recorded a subset of neurons from P12-13 rats in current-clamp mode in the presence of NMDA (AP5, 50 µM), AMPA (CNQX, 20 µM), and GABA-A (gabazine, 1 µM) receptor antagonists. Nicotine application (300 nM to 10 µM) had no effect on resting membrane potential or rheobase (data not shown, *n* = 3 neurons). Nicotine application also did not increase the spike frequency of these neurons (Fig. 3E). These experiments establish that FS neurons in S1 lack direct postsynaptic nicotinic responses across all developmental stages investigated, indicating that nicotinic modulation of these critical inhibitory neurons likely occurs through other mechanisms.

**Figure 3.**
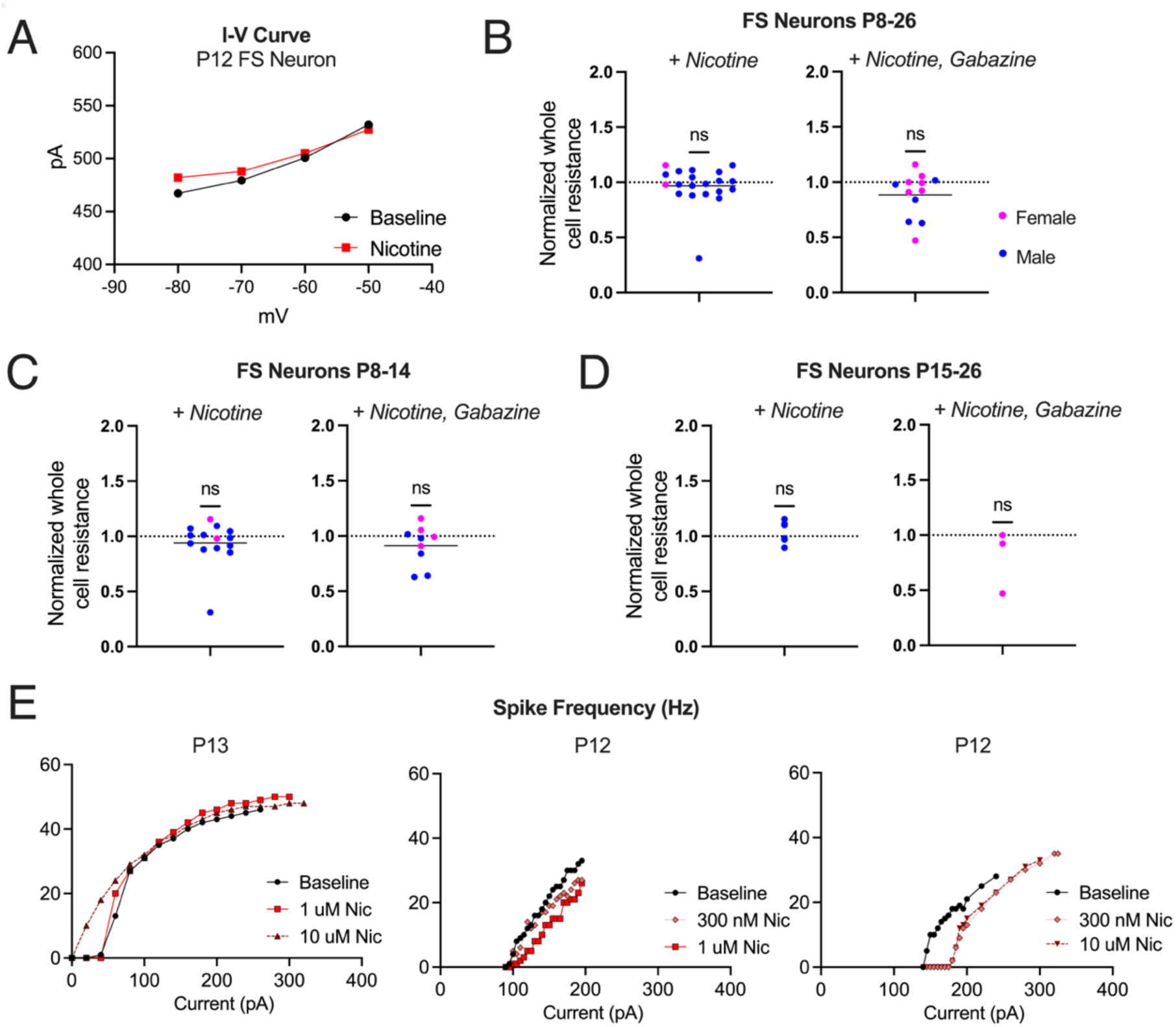
Somatodendritic responses of FS and RS neurons in S1. **A**. I-V curve of a P12 FS neuron before and after nicotine application. **B**. Normalized whole cell resistance of all FS neurons after nicotine application (*left*, *n* = 20, paired *t*-test, t_(19)_ = 0.77, *p* = 0.45) and in the presence of gabazine (*right*, *n* = 12, paired *t*-test, t_(11)_ = 1.81, *p* = 0.10). **C**. Normalized whole cell resistance of FS neurons from P8-14 rats (*left*, *n* = 14, paired *t*-test, t_(13)_ = 0.77, *p* = 0.45) and with gabazine (*right*, *n* = 9, paired *t*-test, t_(8)_ = 1.56, *p* = 0.16). **D**. Normalized whole cell resistance of FS neurons from P15-26 rats (*left*, *n* = 6, paired *t*-test, t_(5)_ = 0.05, *p* = 0.96), and with gabazine (*right*, *n* = 3, paired *t*-test, t_(2)_ = 1.18, *p* = 0.36). **E**. Spike frequency with bath nicotine application with synaptic blockers from 3 FS neurons recorded from rats P12-13.

### Absence of nicotinic responses in FS neurons is not attributable to receptor desensitization

The sustained effects of nicotine bath application (10 µM, 3 to 6 minute application) on inputs to FS neurons (Fig. 2) may represent a non-desensitizing nicotinic mechanism (Askew et. al., 2019). Our finding that bath-applied nicotine did not produce postsynaptic responses in FS neurons (Fig. 3) suggests that rapid receptor desensitization may preclude observation of postsynaptic nicotinic effects on these cells. To test this, we focally applied puffs of nicotine (20 to 30 µM) using a Picospritzer, while recording in current clamp in the presence of synaptic blockers (Fig. 4A-B). Nicotine did not depolarize any FS neuron at any age (Fig 4C, E). In contrast, nicotine depolarized one third of RS neurons by an average of 4.6 mV (Fig. 4D-E), in both younger and older pups. This result is consistent with previous findings that nicotine induces small depolarizations in some RS neurons in sensory cortical areas (Askew et al., 2019), though previous literature has not demonstrated this effect in the presence of synaptic blockers. As this brief nicotine application method reduces the likelihood of receptor desensitization, this suggests that FS neurons lack postsynaptic responses to nicotine. These results demonstrate that FS neurons do not have postsynaptic responses to nicotine applied via bath or directly to the soma, as early as these neurons can be reliably identified through early adolescence.

**Figure 4.**
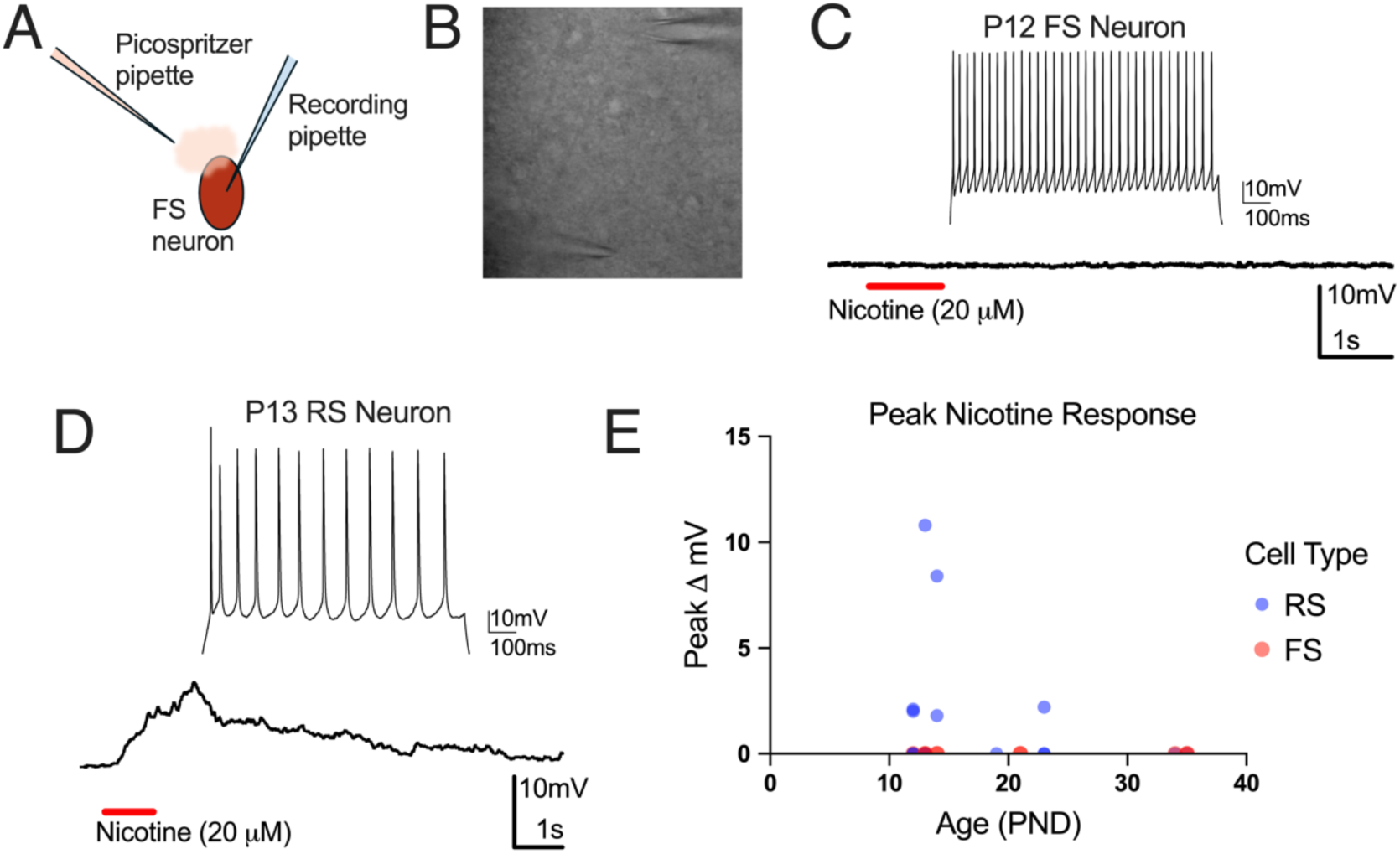
FS & RS responses to direct nicotine application. **A**. Schematic for nicotine puff recordings. **B**. Image of puff recording with recording pipette (*top*) and puff pipette (*bottom*). **C**. Representative recording from a FS neuron with direct nicotine application. **D**. Representative recording from a RS neuron that depolarized in response to direct nicotine application (average of 5 responses). **E**. Summary of voltage changes from RS and FS neurons with direct nicotine application at various ages (*N* = 10 animals, *n* = 10 FS neurons; 18 RS neurons).

### FS neurons express two common nicotinic receptor subunits

The α4β2 heteromeric and α7 homomeric nAChRs are the most commonly expressed nicotinic receptors in mammalian cortex (Millar & Gotti, 2009; Nair & Liu, 2019). Fig. 5A-B depicts images of sections through the barrel cortex of P12 rats, demonstrating localization of mRNA for PV and for CHRNA4 and CHRNA7, the genes coding for the α4 and α7 nicotinic subunits, respectively. At P12, 64.2% of PV+ neurons expressed CHRNA4, and 75.1% expressed CHRNA7 (Fig. 5C). More than half of PV+ neurons (59.8%) at P12 co-expressed mRNA that encode α4 and α7 nAChR subunits (Fig 5C, right). At P19, 59.7% of PV+ neurons expressed α4 nAChR-encoding mRNA, 80.0% expressed α7-nAChR-encoding mRNA, and 45.4% co-expressed both mRNAs (Fig. 5C). These expression levels did not significantly differ between P12 and P19. These data indicate that PV neurons express mRNAs that encode α4 and α7 nAChR subunits from early in development, near the time point when these neurons develop characteristic fast-spiking properties.

**Figure 5.**
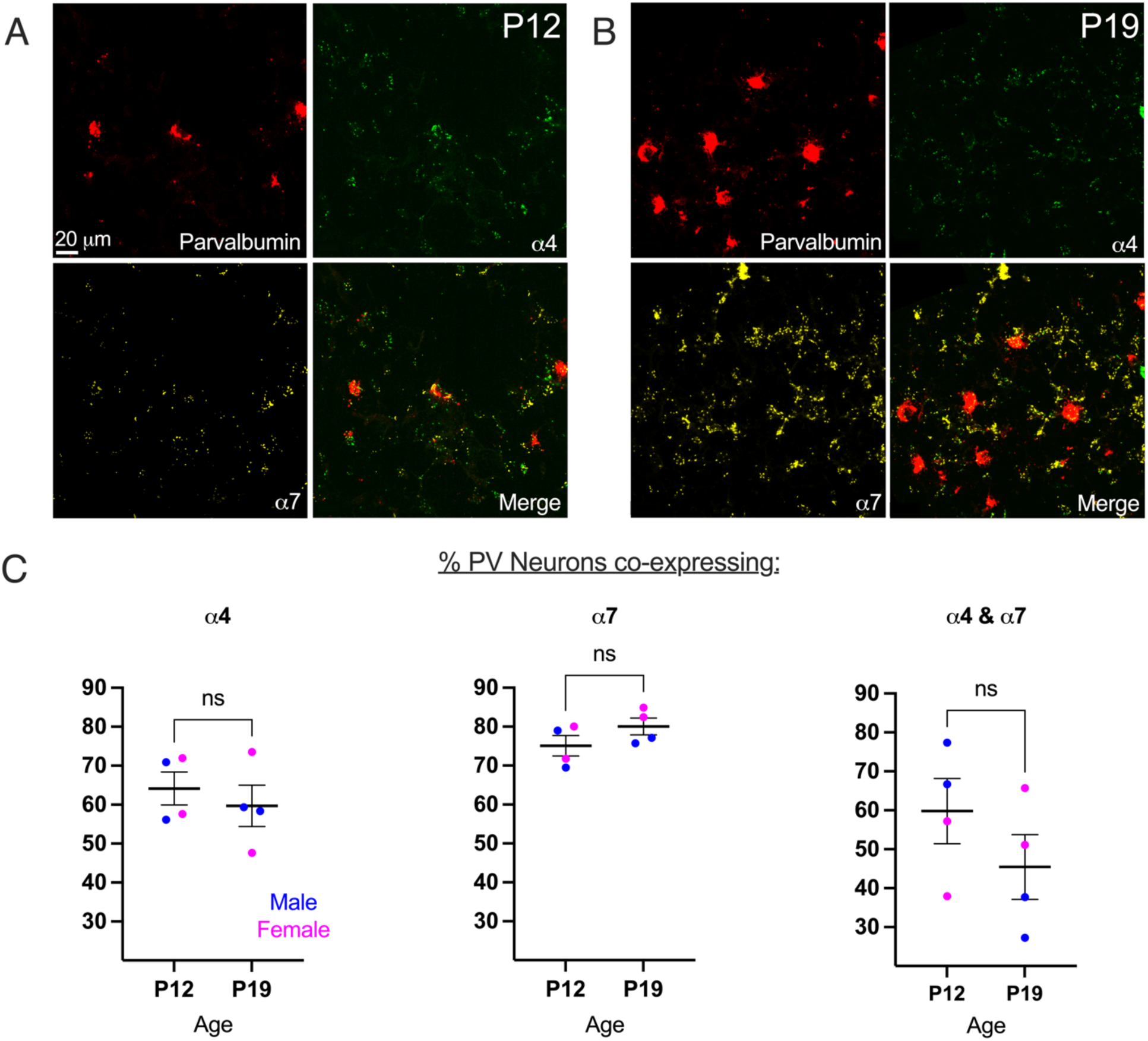
PV neurons express mRNAs that encode multiple nAChR subunits in S1: α4 and α7. RNAscope images from S1 in a P12 (***A***) and P19 (***B***) rat showing co-expression of α4 and α7 mRNA in neurons also expressing PV. **C**. Co-expression levels of nAChR mRNA in PV+ neurons from P12 and P19 rats (*left,* α4: *N* = 2 rats/age, *n* = 2 sections/rat, unpaired *t*-test, t_(6)_ = 0.65, *p* = 0.54; *center,* α7: *N* = 2 rats/age, *n* = 2 sections/rat, unpaired *t*-test, t_(6)_ = 1.46, *p* = 0.20; *right*, α4* & α7: *N* = 2 rats/age, *n* = 2 sections/rat, unpaired *t*-test, t_(6)_ = 1.22, *p* = 0.27). Means ± 95% CI (**C**).

### Lynx1 expression increases early in development in S1

The expression of mRNAs that encode nAChR subunits by PV neurons, including in younger animals, contrasts with the lack of responses of these cells to nicotine. One mechanism that may prevent activation of nAChRs is the expression of lynx1 protein. Lynx1 is a nAChR modulator co-expressed with nAChRs in the brain (Ibañez-Tallon et al., 2002). Lynx1 acts as a molecular brake on nicotinic signaling by orthosterically binding nAChRs, thereby limiting the elevated levels of cortical plasticity observed during developmental critical periods to their more stable, characteristic adult states (Morishita et al., 2010; Miwa et al., 2021; Takesian et al., 2018). Studies of lynx1 control of plasticity have focused on auditory (Takesian et al., 2018) or visual cortex (Morishita et al., 2010; Bukhari et al., 2015; Sadahiro et al., 2016; Sajo et al., 2016). The timeline of lynx1 expression and its role in the somatosensory cortex is not known.

We examined whether lynx1 is expressed in the developing S1 cortex and compared this expression to that in the visual cortex. We used RT-qPCR to quantify *lynx1* mRNA levels in rat S1 and primary visual cortex (V1) across developmental ages. Lynx1 mRNA was not detectable in neonates (P0), underwent a gradual increase in the first postnatal week (P7), and greatly increased in the second postnatal week (Fig. 6A). In V1, lynx1 mRNA expression profile was similar, as the greatest increase in mRNA expression occurred during the second postnatal week (Fig. 6B).

**Figure 6.**
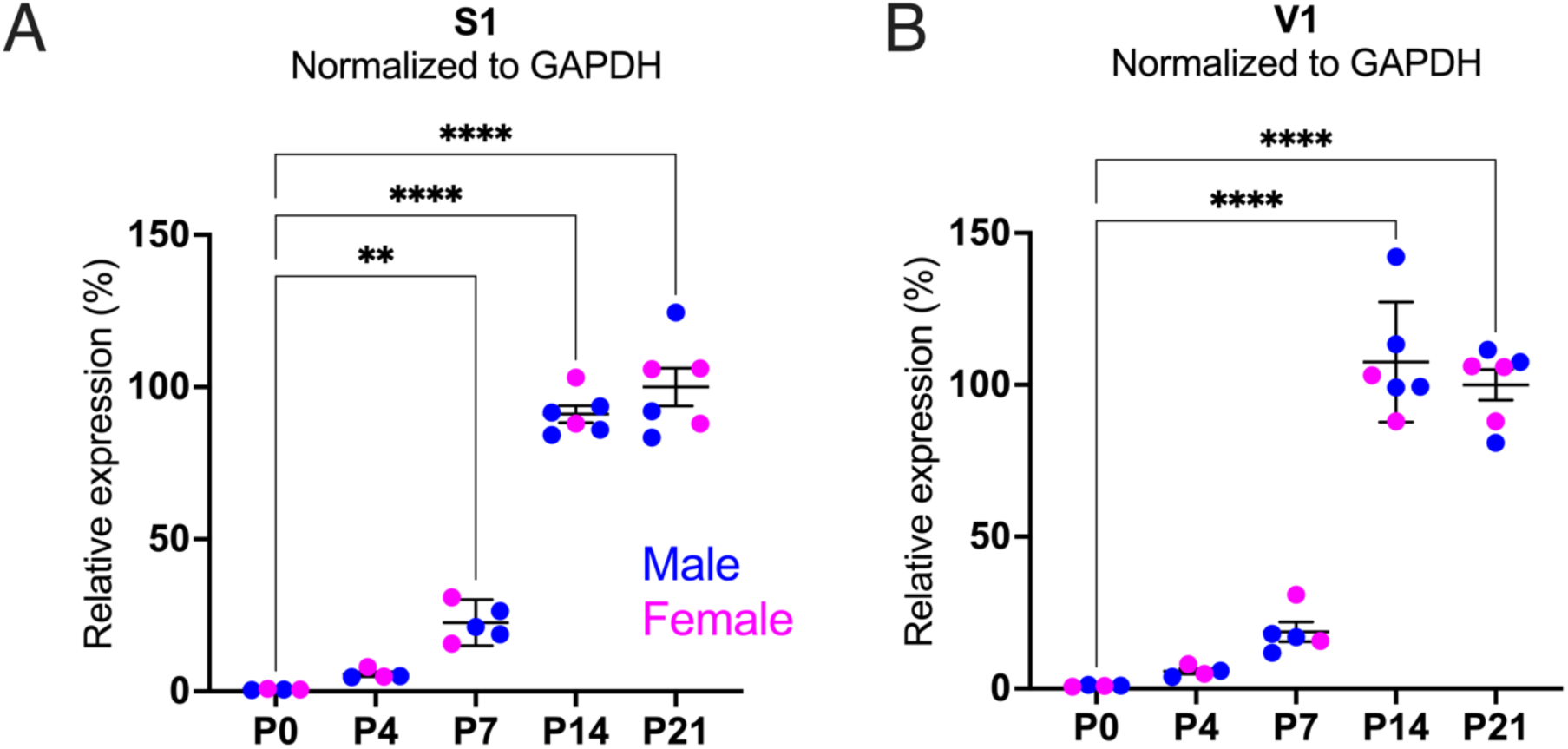
*Lynx1* expression increases in S1 and V1 in the second postnatal week. Relative mRNA expression levels of *lynx1* in S1 (**A**, *n* = 4-8 animals at each age, F(4,20) = 150.4, *p* <0.0001, one-way ANOVA, Dunnett’s multiple comparison tests) and V1 (**B**, *n* = 4-8 animals at each age, F(4,20) = 102.7, *p* <0.0001, one-way ANOVA, Dunnett’s multiple comparison tests). Means ± 95% CI (**A**, **B**)

To examine the expression of *lynx1* mRNA specifically in PV neurons we took advantage of *in situ* hybridization (RNAscope). We studied mouse S1 during the second postnatal week (P12, Fig. 7A) and during adulthood (P100, Fig. 7B). At P12, 96.4% of PV+ neurons expressed *lynx1* (data not shown, *N* = 2 animals per time point), and at P100, 83.5% expressed *lynx1* (data not shown, *N* = 2 animals per time point), confirming that PV neurons express mRNA for a protein that can act as a brake on nicotinic receptor function from nearly as early as they can be reliably identified and recorded using electrophysiology. Co-expression of *lynx1* mRNA remains high in PV neurons well into adulthood.

**Figure 7.**
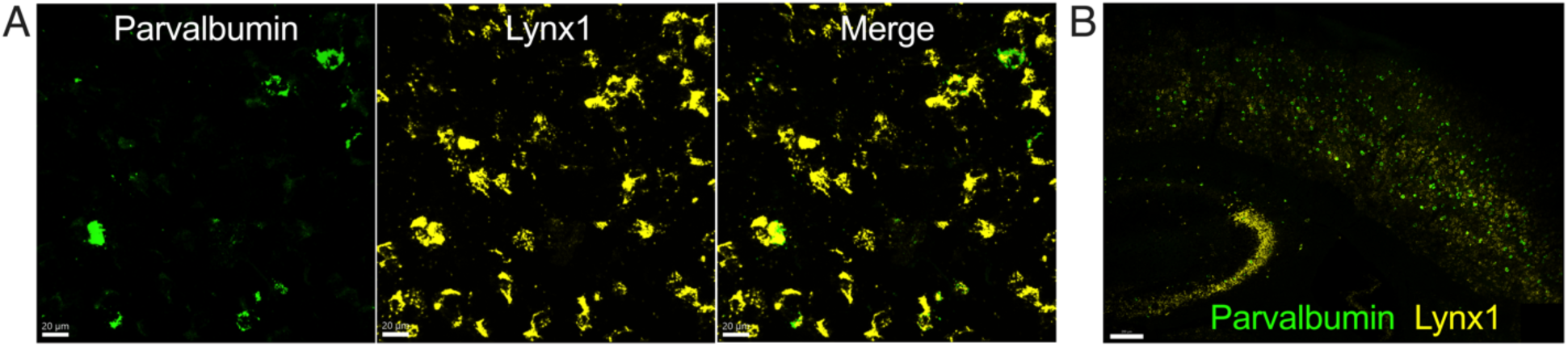
Lynx1-encoding mRNA is expressed in PV neurons in mice early in development through adulthood. RNAscope images from S1 in a P12 (***A***) and P100 (***B***) mouse showing co-expression of *lynx1* mRNA in neurons also expressing parvalbumin. Characteristic *lynx1* expression pattern in adult hippocampus is shown in (***B***). See *Results* for expression levels.

## DISCUSSION

We established that nicotine, acting presynaptically, enhances excitatory inputs to FS neurons as early as the second postnatal week in rats, a period corresponding to late human gestation (Clancy et al., 2007). This is consistent with previous work demonstrating that nicotine or ACh, acting at nAChRs, increase frequency of excitatory inputs to adult cortical FS neurons (Couey et al., 2007; Kassam et al., 2008). The source of these inputs has not been identified, but intrinsic axon collaterals from corticothalamic neurons are a possible component (West et al., 2006; Kassam et al., 2008; Heath et al., 2010; Kim et al., 2014). In the present study, we successfully identified FS neurons at younger ages and found that nicotine increases the frequency of spontaneous excitatory postsynaptic currents (sEPSCs) in 57.1% of FS neurons from as early as P8.

The sensitivity to nicotine of synaptic inputs to FS neurons presents a unique vulnerability to exogenous nicotine exposure, such as from maternal smoking or vaping. These effects may be driven by a less well-characterized nicotinic receptor, such as the ⍺4β2⍺5 receptor expressed in layer 6 pyramidal neurons that exhibits nicotinic responses that are highly resistant to desensitization, similar to the persistent effects on sEPSC frequency in our recordings (Heath et al., 2010; Bailey et al., 2014; Venkatesan et al., 2023). This receptor is implicated in many of the long-term changes resulting from nicotine exposure during prenatal or early postnatal development (Heath et al., 2010), and the CHRNA5 gene coding for the ⍺5 nAChR subunit is linked to nicotine addiction, schizophrenia, and effects of developmental nicotine exposure (Hong et al., 2012; Bailey et al., 2014).

In contrast to the effects on presynaptic inputs, we found no evidence for nicotine having direct effects on the somatodendritic compartment of FS neurons at any time during development. The results in older animals are consistent with previous literature demonstrating that nicotinic agonists do not depolarize FS neurons in adults (Porter et al., 1999; Couey et al., 2007; Gulledge et al., 2007). We now show that FS neurons from even younger animals—during the second postnatal week—do not respond to nicotine.

To determine if the lack of responses is related to the expression—or lack thereof—of nicotinic receptors, we established a developmental timeline of expression of mRNAs encoding the α4 and α7 nAChR subunits, the most commonly expressed in mammalian cortex (Millar & Gotti, 2009; Nair & Liu, 2019), in S1 PV neurons. Both receptors are developmentally regulated (Naeff et al., 1992). We established that, early in development, nearly 2/3 of FS neurons in rat S1 express the gene coding for α4 nAChR subunits, and 3/4 of them express the gene coding for α7 nAChR subunits. More than half express mRNAs encoding both receptor subunits. This reveals a heterogeneity in expression of nAChR subunits that may correspond to different PV neuron subtypes, such as basket and chandelier cells. Very few PV+ neurons expressed neither mRNA-encoding receptor subunit. Despite the known early upregulation of these mRNAs in cortical neurons (Naeff et al., 1992), the expression profile in PV neurons changed little between P12 and P19 in our analysis.

The expression of these nAChR subunits in PV neurons, their early emergence in development, and relatively stable expression contrasts with the lack of responses of these neurons to nicotinic stimulation. We considered possible mechanisms for this contradiction. Whether nAChRs in PV neurons are expressed in somatodendritic compartments or in axon terminals has not been established. Nicotine enhances frequency and amplitude of sIPSCs in cortical pyramidal neurons, possibly through effects on 5HT3AR+ inhibitory neurons (Couey et al., 2007; Takesian et al., 2018; Askew et al., 2019). Further studies may reveal whether α4β2, α7, ⍺4β2⍺5, or other nAChRs play a role in transmitter release at the terminals of FS neurons.

Lynx1, a prototoxin that serves as a brake on nicotinic signaling in other cortical areas (Morishita et al., 2010), is expressed in FS neurons, but its role in these neurons is unclear. Lynx1 is part of the large Ly6/uPAR/neurotoxin superfamily whose members are often expressed alongside nAChRs as accessory proteins (Miwa et al., 2021).

Lynx1 is transported to the cellular membrane, binds to nAChRs and reduces their signaling in a similar manner to toxins in elapid snake venom (Miwa et al., 1999). We revealed a large developmental increase in expression of the mRNA that encodes lynx1 in S1 and confirmed that it is expressed in PV neurons in the second postnatal week.

Changes in the expression of lynx1 across development have been studied only in visual and auditory cortex, and only at later developmental stages (≥ P18). Here, we provide the first analyses of lynx1 expression starting at birth, in both V1 and S1. We find that, in S1, *lynx1* mRNA expression is not detectable in neonates but increases sharply around P7. In V1, *lynx1* mRNA expression profile was similar, as the greatest increase in mRNA expression occurred during the second postnatal week.

The sharp increase in lynx1 expression suggests that, at earlier ages, when S1 is undergoing rapid development, FS neurons may be regulated by nicotinic currents when lynx1 expression is low or absent. Because FS neurons in rats cannot be reliably identified this early (see above), we could not test this hypothesis directly. Developmental changes in lynx1 regulation of nicotinic responses may be involved in shaping plasticity and critical periods in S1 development (Erzurumlu & Gaspar, 2012). In V1, lynx1 expression increases at the end of the critical period for monocular deprivation (Morishita et al., 2010). In mice lacking the *lynx1* gene, this window of plasticity remains open well into adulthood (Morishita et al., 2010). Converging evidence implicates lynx1 in the closure of critical periods across sensory cortex, with a different developmental profile in each (Anderson et al., 2020). Lynx1 may be an important nicotinic plasticity brake in multiple cortical areas, which presents treatment possibilities for perinatal nicotine exposure, as well as other cognitive and developmental disorders that involve impaired cortical plasticity. Its role and mechanism in FS neurons during the first two weeks of development remains to be explored. Nicotinic dysfunction in FS neurons is associated with numerous neuropsychological and neurological conditions, but nicotinic mechanisms driving development of FS neurons are understudied.

